# Diffusion MR Microscopy of the Human Hippocampus at 50 μm Resolution at 14.1 T for 3D Virtual Histology

**DOI:** 10.64898/2026.09.04.749425

**Authors:** Yao Shen, Ziyang Yu, Qinfeng Zhu, Zhiyong Zhao, Pan Wang, Yu Yin, Peiran Jiang, Chengbo Wang, Xueqian Kong, Jing Zhang, Dan Wu

**Affiliations:** Department of Biomedical Engineering, College of Biomedical Engineering & Instrument Science, Zhejiang University, Hangzhou, China; Children’s Hospital, Zhejiang University School of Medicine, National Clinical Research Center for Child Health, Hangzhou, China; Department of Pathology, The First Affiliated Hospital, Zhejiang University School of Medicine, Hangzhou, Zhejiang, China; National Human Brain Bank for Health and Disease, Zhejiang University, Hangzhou, Zhejiang, China; Institute of Translational Medicine SJTU, Shanghai, China; Department of Neurology of the Second Affiliated Hospital and Department of Neurobiology, Zhejiang University School of Medicine, Hangzhou, China; Department of Electrical and Electronic Engineering, Faculty of Science and Engineering, University of Nottingham Ningbo China, Ningbo, Zhejiang, China

**Keywords:** Diffusion MR microscopy, Human hippocampus, 3D virtual histology, Ultrahigh field, Tractography, Structure tensor analysis

## Abstract

Bridging the gap between macroscopic neuroimaging and microscopic histology remains a challenge in neuroscience. Here, we present a technical feasibility study of diffusion MR microscopy (dMRM) in the *ex vivo* human hippocampus using a 14.1 T ultrahigh-field MRI system. Mid-hippocampal blocks from control and Alzheimer’s disease (AD) patients were acquired at 50 × 50 × 100 µm³, representing, to our knowledge, the highest resolution reported for human brain diffusion MRI. At a spatial scale approaching cellular dimensions, dMRM maps and tractography showed close correspondence with immunohistological markers of axons and dendrites. The acquired data delineated six hippocampal laminae, distinguished the inner and outer molecular layers, and preserved short, lamina-specific trajectories that became attenuated or blurred at 100 µm resolution. Tractography further depicted the major components of the hippocampal tri-synaptic circuit—the perforant path, mossy fibers, and Schaffer collaterals in 3D. In the AD specimen, dMRM revealed reduced laminar contrast, local discontinuities, and altered radial and tangential fibers that were paralleled by pathological staining. With the 3D nondestructive characterization of human brain microstructure and circuitry, *ex vivo* dMRM showed its potential toward the goal of 3D virtual histology.

## Introduction

Deciphering the intricate cytoarchitecture and neural circuitry of the human brain is fundamental to understanding cognitive functions and neurodegenerative pathologies. The hippocampus is one of the most complex structures in the human brain, plays a vital role in memory and navigation^[1,2]^, and is selectively vulnerable in several neurological and psychiatric disorders^[3–6]^. Its intricate cytoarchitecture, subfield divisions, laminar organization, and complex interconnections call for three-dimensional (3D) characterization of hippocampal microstructure, connectivity, and pathology at mesoscopic or even microscopic scales.

Among the various approaches available for structural characterization, histological examination remains the gold standard because it provides exquisite microscopic contrast. However, conventional histology is inherently destructive, is typically restricted to two-dimensional (2D) sections, and is susceptible to geometric distortions introduced during tissue cutting, mounting, and staining. Conversely, magnetic resonance imaging (MRI) preserves tissue integrity and provides reproducible, three-dimensional, multiparametric representations of the human brain, but has historically been limited by spatial resolution. While clinical MRI (typically >1 mm) suffices for gross volumetric assessment^[7–10]^, it fails to resolve the mesoscopic and microscopic organization essential for characterizing complex structures. Consequently, there is an urgent need to bridge the substantial resolution gap between *in vivo* MRI at the macroscale and histology at the microscale, aiming for a “3D virtual histology” that combines the benefits of both modalities.

*Ex vivo* MRI at ultrahigh-field (UHF; ≥7 T) has emerged as a powerful tool for bridging this gap. Benefiting from the near-linear gain in signal-to-noise ratio (SNR) with field strength^[11,12]^, along with unlimited scan times for *ex vivo* tissue and specialized hardware, this approach has pushed the boundaries of spatial resolution. Particularly, diffusion MRI (dMRI), which utilizes the restricted water diffusion in biological tissues to map tissue microstructure^[13,14]^ and fiber tracts^[15,16]^, has been extensively used in *ex vivo* studies^[17]^. However, pushing dMRI to microscopic scales is far more challenging than T1- or T2-weighted structural MRI, since dMRI relies on strong diffusion-encoding gradients that attenuate the signal, rendering it intrinsically low in SNR. Supplementary **Table S1** summarizes existing *ex vivo* human dMRI studies performed at UHF, spanning 7T to 16.4T^[18–32]^, where the spatial resolution has largely plateaued around 100 μm isotropic in recent years. At this resolution, hippocampal laminar structures such as the pyramidal cell layer (PCL) and molecular layer (ML) can be clearly delineated^[19,20,22,33]^. Nevertheless, it remains insufficient for resolving cellular-scale microstructures. For instance, the soma of a human pyramidal neuron is approximately 10–50 μm^[34,35]^; glial cells are even smaller (5-10 μm) ^[36]^; and the entire granule cell layer in the hippocampus can locally be as thin as ∼60 μm^[37]^. Therefore, a further enhanced resolution approaching a cellular scale would be ideal for building MRI towards virtual histology.

In this study, we developed a diffusion MR microscopy (dMRM) protocol using a 14.1 T UHF MRI system. Using specialized sample preparation and a 3D volumetric dMRI pulse sequence, we achieved an acquired resolution of 50 × 50 × 100 µm³, corresponding to the highest spatial resolution among the *ex vivo* human diffusion MRI studies summarized in Supplementary Table S1. Whereas previous studies demonstrated that major hippocampal layers and intrahippocampal pathways could be examined at approximately 100–300 µm resolution^[18–20,22,23,25,32]^, the present study aims to show whether pushing the resolution further could improve characterization of thin laminae, laminar subdivisions, and finner tractography that remain vulnerable to partial-volume averaging. We took advantages of structure-tensor analysis of NF and MAP2 immunohistochemistry to answer this question. We further applied the mesoscopic dMRM to an Alzheimer’s disease specimen to illustrate its potential for depicting local microstructural and pathological alterations. Together, this study establishes a technical framework for continuous, 3D, and nondestructive characterization of human hippocampal architecture, circuitry, and pathology, advancing dMRM toward the goal of 3D virtual histology.

## Methods

### Brain tissue preparation

Human brain tissues were obtained from the National Human Brain Bank for Health and Disease of China in Zhejiang University (CNBB). Written informed consent, including an anatomical gift act and repository consent for data sharing, was obtained from all donors or their legal representatives. The diagnosis of AD was confirmed at autopsy, and the control was selected from cases without vascular or other neurological complications. The study included one subject without known brain pathology and one subject with AD; their specific information was summarized in Supplementary **Table S2**.

The hemisphere was fixed in 4% paraformaldehyde for at least 30 days, and then the middle part of the right hippocampus was dissected carefully from the coronal brain section (approximately 5 mm thick). Before MRI, tissue specimens were incubated in phosphate-buffered saline (PBS) containing 1 mM gadolinium-based contrast agent (Gd-DTPA, Berlex Imaging, Wayne, NJ, USA) for 7 days at 4°C to enhance MR signals. The samples were transferred to 50 mL polypropylene centrifuge tubes with supporting materials to prevent movement during scanning. Finally, the tubes were filled with Fomblin (Solvay Solexis, Thorofare, NJ, USA) to prevent dehydration and susceptibility artifacts.

### MRI data acquisition

Ultrahigh-resolution *ex vivo* dMRI was performed on a 14.1T Bruker Avance III HD NMR instrument (Bruker Biospin, Billerica, MA, USA) equipped with a micro 2.5 gradient coil (maximum gradient strength = 1.5 T/m and slew rate =15,000 T/m/s) and a 30 mm diameter volume transceiver. A 3D echo planar imaging (EPI) sequence was applied with the following parameters: repetition time = 1000 ms, echo time = 24 ms, field of view(FOV) = 16 × 19 × 5 mm^3^, resolution = 50 × 50 × 100 µm^3^, b-value = 3000 s/mm², 20 diffusion directions uniformly distributed in space, and 2 non-diffusion-weighted measurements. Each diffusion-weighted volume was acquired with 16 averages, resulting in a total acquisition time of 78 hours per sample.

This protocol was carefully optimized to balance spatial resolution, angular resolution, SNR, and total scan time. Pushing spatial resolution to the microscopic scale (voxel volume = 2.5 × 10⁻⁴ mm³) imposes severe penalties on both SNR and acquisition time. Our pilot experiment demonstrated that an acquisition scheme with fewer averages (e.g., 10 averages with 10 directions) caused the raw SNR of b=3000 s/mm^2^ images to drop far below the recommended threshold of 10 ^[22]^, leading to significant bias in tensor estimation, as evidenced by the noisy mean diffusivity (MD) map (**Fig. S1**). Therefore, within the practical constraint of total scan time, increasing the number of signal averages was prioritized over further increasing the number of diffusion directions.

### dMRI data processing

#### Preprocessing and reconstruction

dMRI data were processed using MRtrix3^[38]^. Preprocessing steps included denoising, followed by eddy current and field bias correction. Motion correction was not applied, as no significant movement was observed between scans. Diffusion Tensor Imaging (DTI) was reconstructed by performing an eigenvector analysis on the calculated tensor. Subsequently, fractional anisotropy (FA), mean diffusivity (MD), radial diffusivity (RD), axial diffusivity (AD), and color-coded FA (colorFA) maps were computed. Moreover, Fiber Orientation Distribution (FOD) was estimated using the constrained spherical deconvolution (CSD) reconstruction algorithm^[39]^. The maximum spherical harmonic degree was set to 4 (*l_max_*=4), which mathematically requires at least 15 unique diffusion directions^[39,40]^.

#### Tractography

Fiber tracking of the specimen was performed within MRtrix3, with the following parameters: one seed per voxel with random positioning, a FA threshold of 0.02, an angular threshold of 60°, and a step size equal to 1/10 of the voxel size (5 µm for a 50 µm resolution). The minimum and maximum tract length were set to 0.1 mm (twice the voxel size) and 20 mm, respectively. A tensor probabilistic tracking algorithm was employed^[41]^.

### Immunohistochemistry

After MRI scans, both hippocampi were paraffin-embedded, sectioned at 10 μm, and stained with neurofilament (NF), Microtubule-associated protein 2 (MAP2), hyperphosphorylated tau (p-tau), amyloid-β (Aβ), and counterstained with hematoxylin.

Immunohistochemical staining was performed as follows. Tissue sections were deparaffinized, rehydrated, and subjected to antigen retrieval using a citrate buffer. To quench endogenous peroxidase activity, the sections were incubated with a 0.3% hydrogen peroxide solution for 15 minutes at room temperature. After washing with 1× PBS, the sections were blocked with a solution containing 10% normal goat serum for 1 hour at room temperature to reduce nonspecific background. The sections were then incubated overnight at 4°C with anti-MAP2 (1:200 dilution, Abcam, ab254264), anti-68kDa Neurofilament/NF-L (1:200 dilution, Abcam, ab223343), β-amyloid 1-42 (1:200 dilution, Affinity, AF0019), and phospho-tau (1:200 dilution, Thermo, Thr217) Polyclonal antibodies. Following extensive washing with PBS, the sections were incubated with horseradish peroxidase (HRP)-conjugated secondary antibodies for 1 hour at room temperature. Immunoreactivity was visualized using a 3,3’-diaminobenzidine (DAB) substrate kit, and the nuclei were counterstained with hematoxylin. Finally, the sections were dehydrated, cleared in xylene, and mounted with a resin-based mounting medium. Images were acquired under bright-field illumination using an Olympus VS200 slide scanner (Olympus, Japan) at 20× magnification.

### Structure tensor analysis

Structural orientation in the histological sections was analyzed using structure tensor analysis^[42,43]^ in ImageJ (**Fig. S2**). Briefly, the original immunohistochemical image was first decomposed into three channels using the color deconvolution tool: the first channel represented hematoxylin-stained nuclei, the second channel contained the positive signal of interest (e.g., MAP2-labeled dendrites), and the third channel corresponded to the background. The second channel was then converted into an 8-bit grayscale image. Subsequently, structure tensor analysis was performed using the OrientationJ plugin, which generated an HSB-formatted map of local fiber orientation, where hue corresponds to orientation, saturation to anisotropy, and brightness to the original signal intensity. The structure tensor was computed from the spatial gradient of image intensity, allowing estimation of local orientation and degree of anisotropy based on the eigenvectors and eigenvalues of the gradient tensor ^[43]^.

## Results

### Diffusion MR microscopy protocol and multi-modal dataset of the human hippocampus

We first acquired ultrahigh-resolution dMRM data of the human hippocampus at 14.1T. Given that the mid-hippocampus contains the typical laminar and subfield organization and is most commonly studied^[19,22,23,32]^, the current study focused on the mid-hippocampal tissue blocks. We utilized a 3D diffusion-weighted echo-planar imaging sequence to reach a resolution of 50 × 50 × 100 µm^3^ that was further interpolated to 50 µm isotropic for tractography. The scanned tissue block was subsequently paraffin-embedded and sectioned for targeted immunohistochemical staining for neuronal and pathological markers (**Fig. 1A**). We performed DTI, FOD reconstruction, and tractography analysis with the dMRM data. For histology, adjacent sections were stained for axons (Neurofilament, NF), dendrites (Microtubule-associated protein 2, MAP2), and Alzheimer’s disease pathology markers, including phosphorylated tau (p-tau) and amyloid-β (Aβ) (**Fig. 1B**).

**Figure 1.**
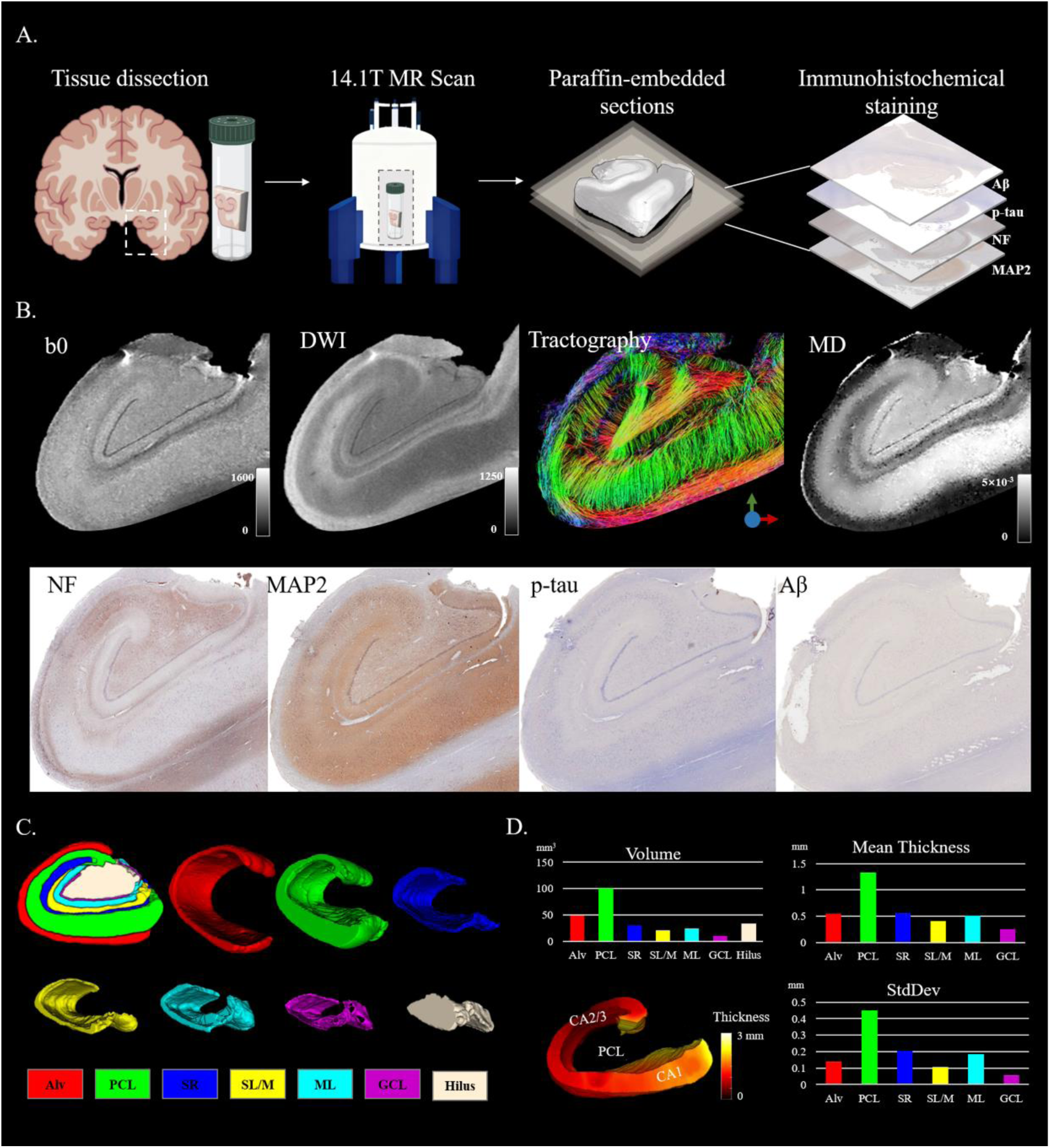
Experimental design and multimodal data. (A) Illustration of the sample preparation, MRI scan, and immunohistochemical staining protocol used in this study. (B) Diffusion MR microscopy (dMRM) and histopathology of a normal human hippocampal specimen (female, 50 years old). dMRM contrasts include non-diffusion-weighted image (b0), diffusion weighted image (DWI) at b=3000s/mm^2^, tractography, and mean diffusivity (MD) maps. Histopathology includes Neurofilament (NF), Microtubule-associated protein 2(MAP2), hyperphosphorylated tau (p-tau) and amyloid-β (Aβ). (C) Manual segmentation of the hippocampus and layer-based quantification of morphological and dMRM features. The layers are labeled as follows: Alveus (Alv), Pyramidal cell layer (PCL), Stratum radiatum (SR), Stratum lacunosum/moleculare (SL/M), Molecular layer (ML), Granule cell layer (GCL) and hilus. (D) Morphological measures of the hippocampal layers.

Leveraging the rich tissue contrast provided by dMRM, we manually segmented the hippocampal layers, including the alveus (Alv), pyramidal cell layer (PCL), stratum radiatum (SR), stratum lacunosum/moleculare (SL/M), molecular layer (ML), granule cell layer (GCL), and hilus (**Fig. 1C**). Specifically, the hippocampal CA subfields were segmented into the PCL, SR, and SL/M layers, whereas the dentate gyrus (DG) was divided into the ML, GCL, and hilus. Furthermore, ML could be further subdivided into the inner molecular layer (IML) and outer molecular layer (OML) in the diffusion-weighted images (DWI) (**Fig. 2B**). Morphologically, the PCL exhibited the largest volume and mean thickness, with the greatest heterogeneity— thickest in CA1 and thinner in CA3/2 (**Fig. 1D**). The alveus, as a white matter fiber layer, showed the second largest volume and thickness, while the GCL was the smallest and thinnest layer (approximately 0.2 mm).

**Figure 2.**
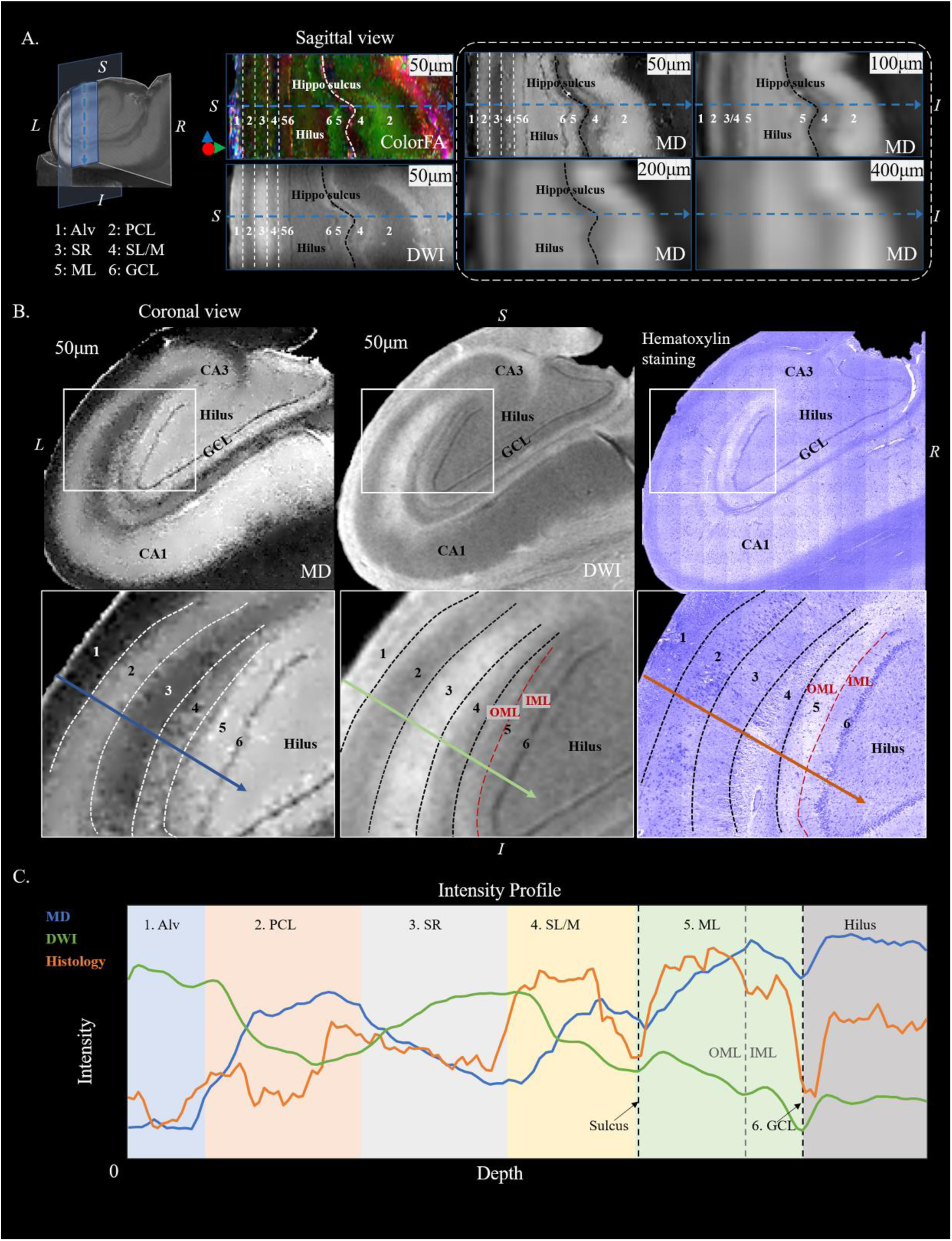
Laminar organization of a normal human hippocampus revealed by dMRM at varying resolutions. (A) Sagittal views of diffusion MRI acquired at 50 µm resolution and subsequently downsampled to 100, 200, and 400 µm isotropic resolutions. At 50 µm, colorFA, DWI, and MD maps clearly delineated six hippocampal layers. At 100 µm, the boundaries between layers became blurred, and the GCL disappeared. At resolutions coarser than 100 µm, these laminar structures could no longer be reliably distinguished. (B) Coronal view at 50 µm resolution alongside hematoxylin-stained histology. (C) Intensity profiles of the MD, DWI and histological maps.

### Diffusion MR microscopy revealed layer-specific microstructural organization of the human hippocampus

Ultrahigh-resolution *ex vivo* dMRM depicted the intricate laminar organization of the human hippocampus (**Fig. 2A**). At a resolution of 50 µm, the layer boundaries were sharply defined. After downsampling to 100 µm isotropic resolution, the layers appeared blurred, e.g., the boundary between the SR and SL/M became blurred, and the thinnest GCL was markedly attenuated. At 400 µm resolution, most laminar structures were no longer distinguishable, leaving mainly coarse gray–white matter contrast.

The coronal view at 50 µm resolution further showed that the DWI and MD maps exhibited complementary contrasts and achieved laminar differentiation comparable to that of histological staining (**Fig. 2B**). Layer-specific intensity profiles were quantified in **Fig. 2C**. In layer 1 (Alv), water diffusion was strongly restricted by dense white matter fibers, resulting in the lowest MD and highest DWI signal. In contrast, layer 2 (PCL) showed high MD and a low DWI signal, corresponding to the densely packed pyramidal neuron soma visible in hematoxylin–eosin staining. Layer 3 (SR) was composed primarily of pyramidal dendrites and Schaffer collateral fibers and exhibited intermediate contrast between layers 1 and 2. Layer 4 (SL/M) contained dendrites and perforant path fibers, and thus showed contrast similar to that of the ML, but with slightly lower diffusivity. Layer 5 (ML), as part of the DG subregion, was mainly composed of granule cell dendrites and displayed higher MD than adjacent regions. Layer 6 (GCL) corresponded to the densely packed granule cell layer, exhibiting strongly restricted diffusion, consistent with the histological findings.

Specifically, in the DWI maps, the ML could be further subdivided into the IML and OML. The IML exhibited lower signal intensity, corresponding to the darker hematoxylin staining (**Fig. 2B and C**). It has been reported that the IML receives associational and commissural inputs from hilar mossy cells, whereas the OML receives perforant path projections from the entorhinal cortex, reflecting a clear spatial segregation of functional inputs^[44]^. Consistent with this organization, tractography with the OML as the termination region showed a prominent bundle of perforant path fibers from the entorhinal cortex (**Fig. S3**).

### Hippocampal neuronal projections revealed by tractography and immunohistochemical staining

Tractography results were also strongly influenced by spatial resolution. As voxel size increased from 50 to 400 µm, the overall streamline density decreased markedly (**Fig. 3A**). Short streamlines ranging from 0.1 to 10 mm were preferentially reduced at 100 µm, whereas longer trajectories, including mossy fibers and the alveus, were relatively preserved. At 200 and 400 µm, both short and long trajectories decreased substantially. These observations indicate that finer spatial sampling can improve the depiction of short-range, layer-specific orientation patterns in the hippocampus.

**Figure 3.**
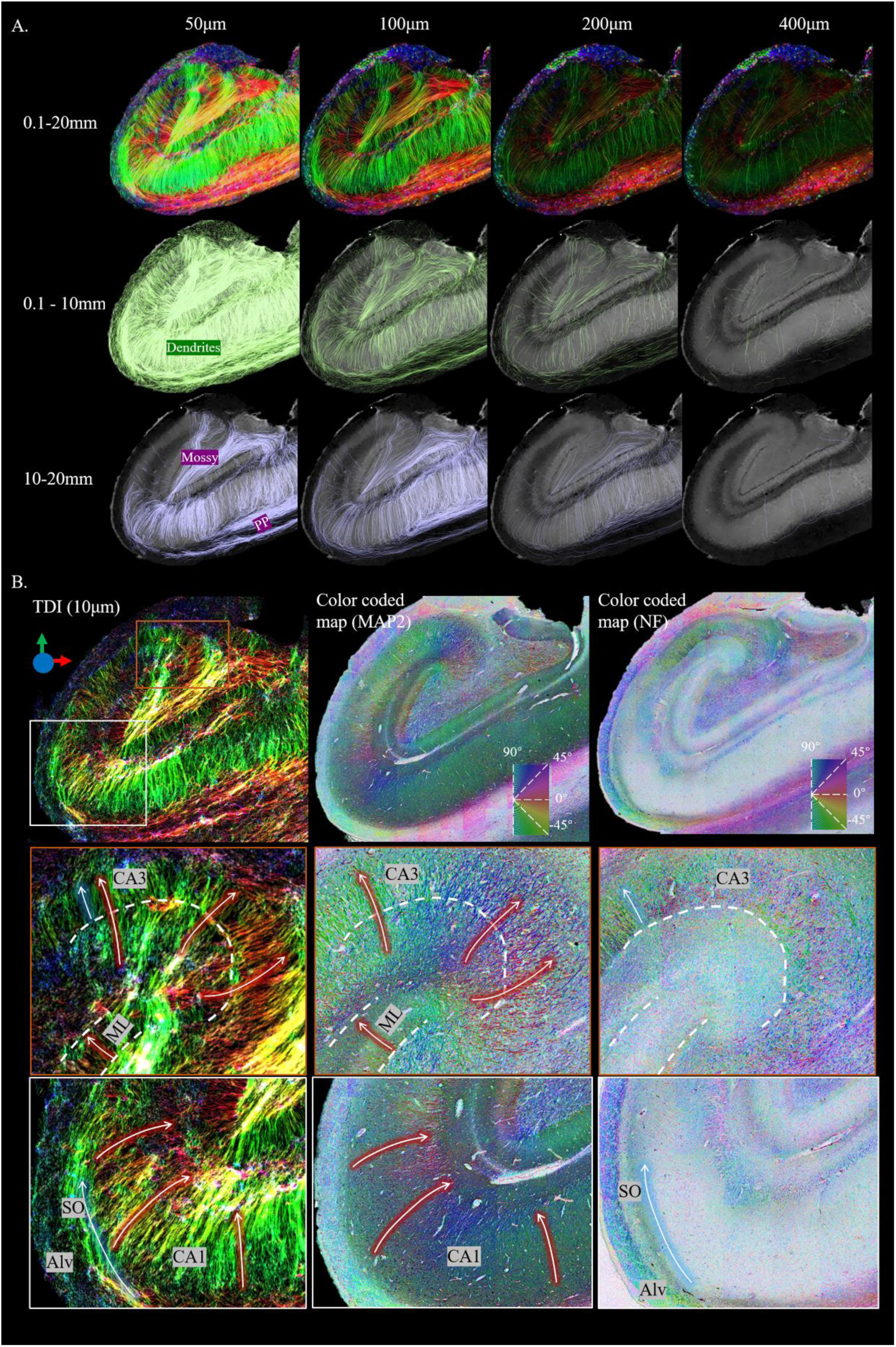
Tractography of the human hippocampus at multiple resolutions and super-resolution TDI. (A) Both short (0.1–10 mm) and long (10–20 mm) fiber populations became sparser as spatial resolution decreased, with short fibers being more severely affected. (B) Super-resolution TDI at 10 μm isotropic resolution provided a level of structural detail comparable to dendritic and axonal histological staining. Glowing red arrows indicate dendritic projections, while glowing blue arrows denote axonal pathways. The orange box highlights the CA3 region, and the white box zooms into the CA1 area. Color coded maps of MAP2 and NF were obtained by structure-tensor analysis of the immunostaining.

To enhance visualization, we performed super-resolution tract density imaging (TDI) reconstruction^[45]^ of the fiber tracts. Based on the 50 µm tractography results, TDI maps were displayed on a 10 µm isotropic grid, achieving a level of detail comparable to that of histology. The resulting TDI maps closely matched the color-coded orientation maps derived from structure-tensor analysis of MAP2 (which labels for dendrites) and Neurofilament (which labels for axons) (**Fig. 3B**). Radial patterns in the CA and ML regions corresponded mainly to MAP2-positive dendrites, whereas patterns in the alveus and SL/M corresponded more closely to NF-positive axonal structures. In CA3 (orange box), where dendritic and axonal signals coexisted, the TDI map likely reflected contributions from both tissue components. In CA1 (white box), fiber trajectories converging from the stratum oriens (SO) into the alveus were validated by NF axonal staining. Similarly, mossy fibers in the hilus, which are known to be unmyelinated axons^[46]^, were also confirmed by NF-positive labeling (**Fig. 3B and 4C**).

**Figure 4.**
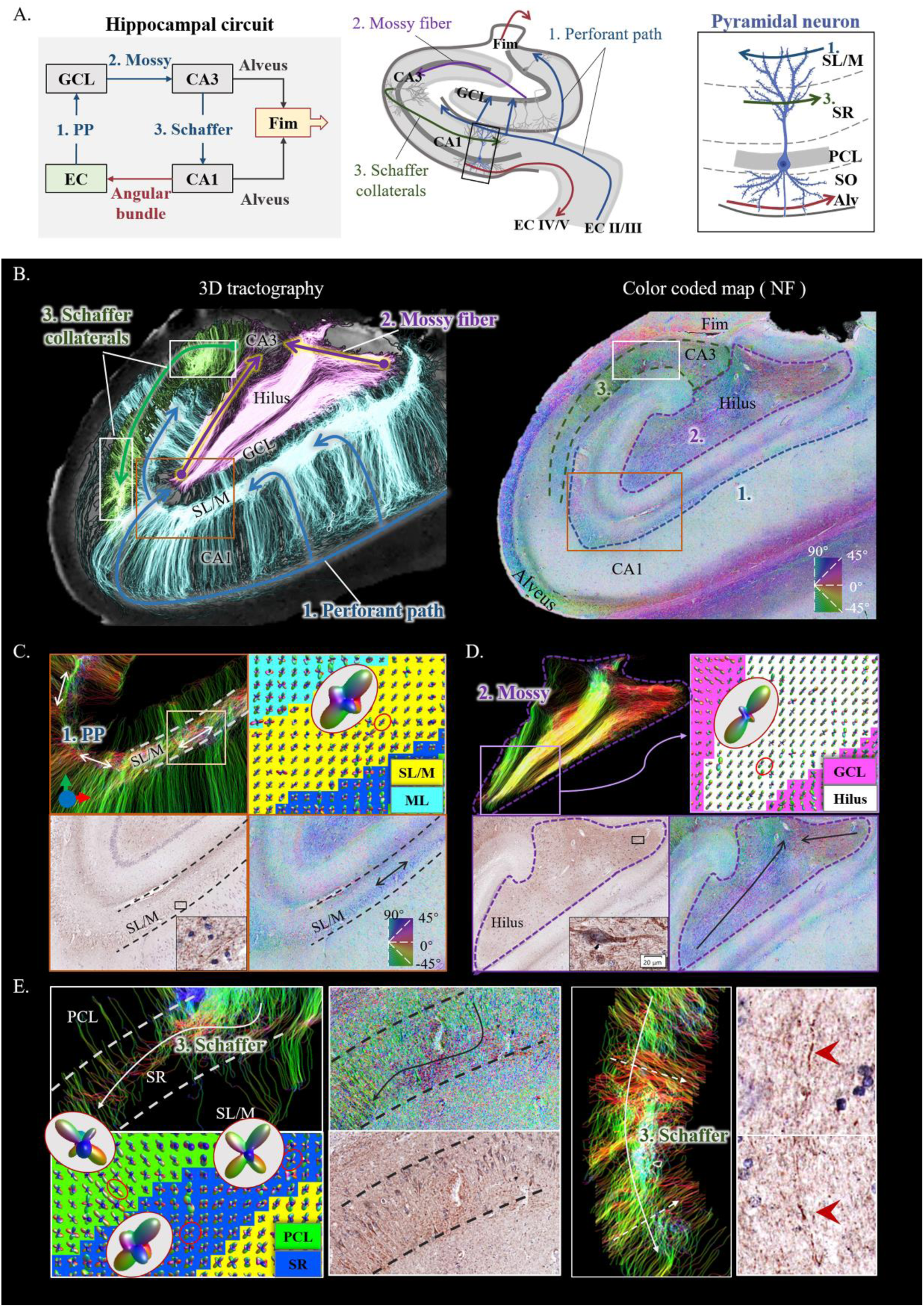
Tri-synaptic pathway reconstructed by 3D tractography. (A) Schematic of the classic hippocampal circuit, illustrating both afferent and efferent pathways. (B) Both 3D tractography and the corresponding NF-stained histological maps delineated the tri-synaptic pathway. The orange region highlights the perforant path (C), the purple region emphasizes the mossy fiber tract(D), and the white region focuses on the Schaffer collateral pathway(E).

Beyond 2D visualization, 3D tractography delineated trajectory groups corresponding to the three components of the classic tri-synaptic hippocampal pathway. This pathway consists of: (1) the perforant path (PP), which projects from the entorhinal cortex to the DG dendrites in the ML, (2) mossy fibers projecting from the DG to CA3, and (3) Schaffer collaterals extending from CA3 to CA1 within the SR layer^[47–49]^ (**Fig. 4A**). Among these, the PP represents the major input to the hippocampus, whereas the primary outputs are conveyed through the fimbria to fornix and the angular bundle projecting back to the entorhinal cortex. With 3D tractography, we were able to identify all three connections of the tri-synaptic pathway, which were further validated by axonal staining (**Fig. 4B-E**).

Specifically, the PP entered the hippocampus tangentially and traversed the CA region, forming synaptic connections with dendrites in the ML and propagating tangentially along the SL/M layer toward CA3. The FOD reconstruction revealed dominant tangential orientations that corresponded to NF staining and its color coded maps (**Fig. 4C)**. The mossy fiber pathway (**Fig. 4D**) was distinctly visible, projecting from the DG to CA3 with excellent spatial agreement among the tractography, FOD, and NF staining results. The Schaffer collaterals originated from CA3 pyramidal neuron axons and projected to CA1, connecting with dendrites in the SR layer. **Fig. 4E** illustrated the CA3 and CA1 regions, and the tangential tracts within the SR-layer corresponding to Schaffer collaterals were clearly resolved. These trajectories were delineated with clearer laminar confinement than is typically available at the 100–300 µm mesoscale^[18,22]^. In addition to these pathways, we also identified fiber bundles within the CA3/2 region oriented along the longitudinal (anterior–posterior) hippocampal axis (**Fig. S4**).

### Diffusion MR microscopy revealed disrupted laminar architecture and dendritic organization in Alzheimer’s disease

To further explore the potential of dMRM for depicting neuropathological alterations, we imaged one *ex vivo* hippocampal specimen from a 91-year-old male donor with AD. Extracellular Aβ plaques were observed in the cortex, whereas intraneuronal p-tau pathology was present in the CA regions (**Fig. 5A**). The AD specimen showed discontinuities in the SR and SL/M layers (as indicated by the triangle in **Fig. 5B**), and reduced interlayer contrast on the MD map. NF staining showed corresponding local structural alterations. Mossy fibers and radial PCL trajectories remained visible (**Fig. 5B and Fig. S5**), while the CA1 PCL appeared thinner than that in the control specimen. MAP2 staining showed a similar local pattern.

**Figure 5.**
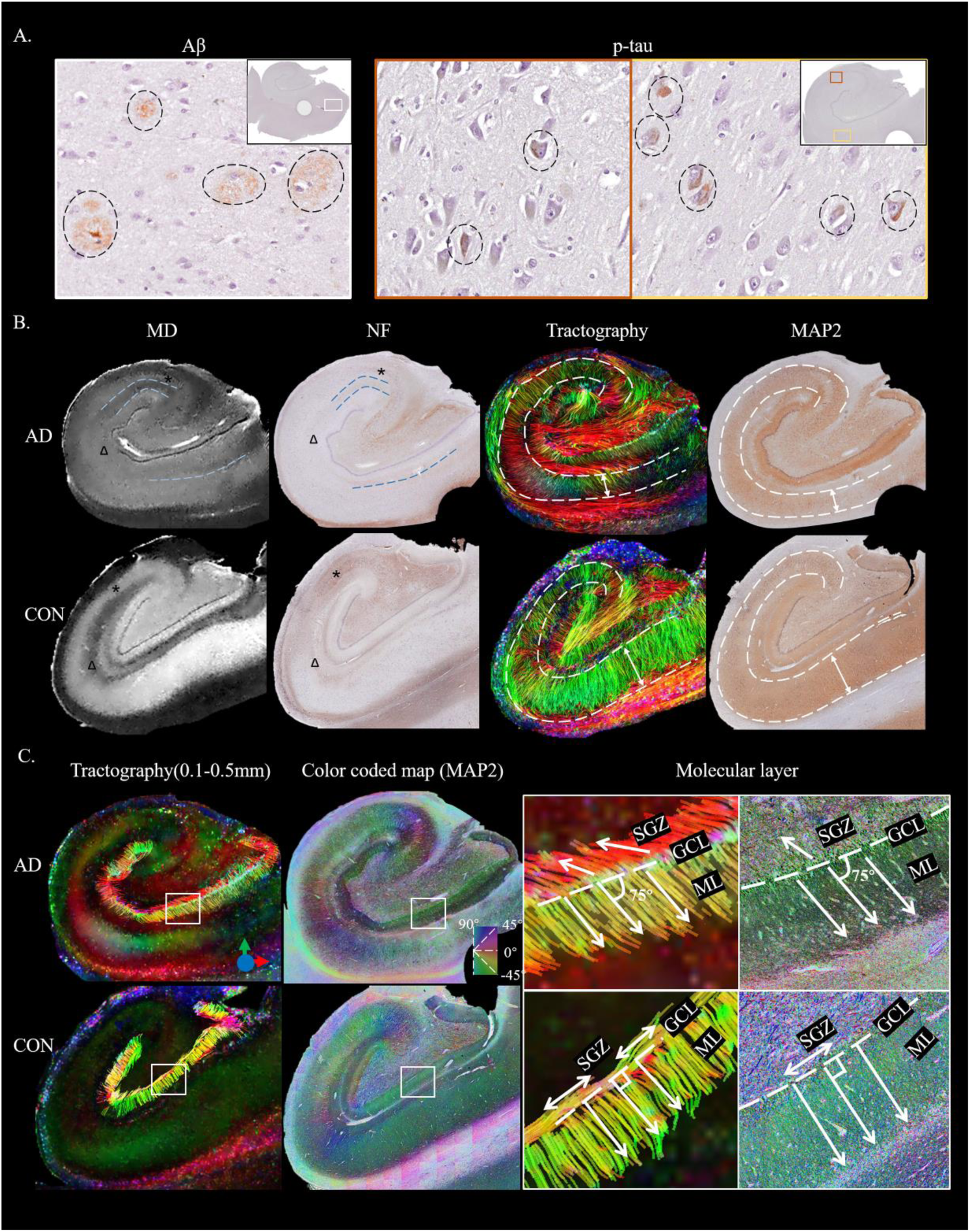
Microscopic laminar and orientation alterations observed in an Alzheimer’s disease specimen. (A) Aβ plaques and p-tau pathology were observed in cortical and hippocampal regions. (B) Laminar architecture and microstructural integrity were markedly disrupted in the AD hippocampus, with reduced interlayer contrast and thinning of the PCL (indicated by white dashed lines and asterisks). Disrupted or missing layers are highlighted with triangles and blue dashed lines. (C) Tractography and MAP2 staining revealed disorganized radial dendritic orientations within the ML and disturbed tangential connectivity within the SGZ in AD.

Using the GCL as the seed region and constraining streamline length to 0.1–0.5 mm, tractography depicted short radial trajectories within the ML that were spatially consistent with MAP2-positive granule-cell dendritic organization (**Fig. 5C**). In the AD specimen, these trajectories appeared less regularly organized and deviated from the predominantly radial arrangement observed in the control specimen. Tangential trajectories near the subgranular zone were also attenuated or displaced. The orientation-encoded MAP2 staining showed qualitatively similar patterns.

## Discussion

Unlike histological staining and optical imaging, MRI provides nondestructive and 3D stereo characterization of biological tissues, but has long been constrained by the trade-off between spatial resolution and SNR. This challenge is particularly acute for diffusion MRI, which is intrinsically SNR-starved due to diffusion-weighted signal attenuation. In this study, we achieved an unprecedented dMRM resolution of 50 µm in the *ex vivo* human hippocampus using a 14.1 T ultrahigh-field scanner, offering a level of detail not previously reported. While previous attempts to achieve 3D histology relied on co-registering serial histological sections to MRI volumes, this process is extremely labor-intensive and prone to geometric distortions from tissue cutting and mounting^[50–52]^, which is difficult for cohort studies. Here, we demonstrated that the ultrahigh-resolution dMRM maps faithfully reflected underlying neural architecture by comprehensive comparison with immunohistology. Consequently, our work establishes a robust technological framework for “3D Virtual Histology” for 3D continuous, nondestructive, distortion-free characterization of tissue microstructure, neuronal projections, and pathology.

Previous ultrahigh-field ex vivo dMRI studies demonstrated that major hippocampal layers could be distinguished at approximately 100–300 µm resolution and that intrahippocampal pathways could be inferred using tractography^[18–20,22,23,25,32]^. The present results extend these mesoscale observations toward finer laminar and circuit-level characterization. At the acquired 50 µm resolution, the data preserved thin laminae, enabled subdivision of the ML into the IML and OML, and depicted short-range trajectories with little partial-volume effect. After downsampling to 100 µm, the SR–SL/M boundary became blurred, the thin GCL was markedly attenuated, and short streamlines were preferentially lost. In particular, the perforant path and Schaffer collaterals were simultaneously delineated with greater laminar specificity than in previous *ex vivo* tractography studies^[18,22,53]^. Combining these observations with NF- and MAP2-derived orientation maps, these findings indicate that the primary benefit of ultrahigh spatial resolution lies not merely in improved anatomical visibility, but also in more accurate characterization of hippocampal laminar organization and local microcircuitry.

Achieving such ultrahigh spatial resolution within a feasible scan time required a deliberate methodological compromise in diffusion acquisition. In the present study, we acquired 20 diffusion directions with 16 signal averages per direction. Sacrificing angular resolution for increased signal averaging is supported by recent mesoscale *ex vivo* studies indicating that once adequate SNR is achieved, the number of diffusion gradient directions has minimal impact on scalar indices and streamline density; indeed, as few as 12 directions have proven sufficient for reliable streamline estimation^[22]^. Importantly, unlike conventional macroscopic imaging that relies on high angular resolution to resolve crossing fibers within large voxels^[54]^, our ultrahigh spatial resolution physically minimizes intra-voxel orientational heterogeneity. By inherently mitigates partial volume effects, principal diffusion orientation can be estimated reliably even with a moderate number of directions^[18,21]^. Thus, the increased spatial resolution effectively compensates for the relatively limited angular resolution.

At the microscopic spatial scale achieved here, dMRM not only revealed distinct laminar patterns but also characterized the underlying cellular environment. Regions rich in glial cells and myelinated fibers (e.g., the alveus and SL/M) exhibited low MD, whereas neuron- and dendrite-dense regions (e.g., PCL and ML) showed higher MD, giving rise to the strong contrast between layers. Notably, although both the GCL and PCL were densely populated with cell bodies, the GCL displayed much lower MD. This discrepancy likely reflected their distinct cellular morphology: pyramidal neurons in the PCL were approximately ten times larger in diameter than granule cells, and the GCL exhibited a much higher packing density (**Fig. S6**), leading to the much more restricted diffusion in the GCL. Moreover, by differentiating the IML and OML, tractography revealed spatial segregation between entorhinal perforant path fibers and commissural/associational fibers from hilar mossy cells. These findings suggested that ultrahigh-resolution dMRM not only identified anatomical layers but also captured functionally distinct microcircuit organization.

The AD specimen illustrates a potential neuropathological application of the protocol. We observed reduced laminar contrast, local discontinuities in the SR and SL/M, and less regular radial and tangential trajectories. Immunohistochemical findings closely paralleled the tractography results. This capability opens the possibility for systematic evaluation of neurodegenerative processes, such as tracking the spatial propagation of Aβ/tau-pathology along neuronal projections^[55–57]^.

This study has several limitations. First, due to tissue fixation and preservation, *ex vivo* MRI is known to differ from *in vivo* MRI^[58]^. Although these biases can be potentially measured and corrected, future studies are needed to bridge the *ex vivo* findings with clinical MRI. Second, as a methodological feasibility study, the current results are based on a limited sample size comprising only one normal and one AD mid-hippocampal block. Although expanding to a larger cohort is necessary for future studies, these representative samples sufficiently highlight the critical benefits of ultrahigh resolution in resolving complex hippocampal pathways and detecting AD-related pathological alterations. Third, while 2D histology was used as the gold standard for validating 3D MRI, it is inherently limited— fibers traversing the plane are truncated, suggesting that future integration with 3D polarized light imaging^[59,60]^ or 3D microscopy^[61,62]^ could provide more comprehensive validation.

Looking ahead, ultrahigh-resolution multimodal MRI may provide increasingly detailed representations of tissue microstructure and serve as a high-resolution reference for interpreting conventional, lower-resolution imaging. In combination with deep-learning approaches, such datasets may enable capturing histological features and predicting selected histopathological patterns from MRI. Continued advances in MRI hardware, acquisition efficiency, reconstruction, and microstructural modeling will be required to translate the *ex vivo* framework to *in vivo* human brain MRI.

## Acknowledgement

This work was supported by the Ministry of Science and Technology of the People’s Republic of China (2021ZD0200202), and National Natural Science Foundation of China (32427802, U24A200313).

## Consent statement

All experiments related to human brain tissues in this study were approved by the Human Ethics Committee of the School of Medicine, Zhejiang University. Written informed consent was obtained from all participants or their legal guardians.

## Reference

1. Burgess N, Maguire E A, O’Keefe J. The human hippocampus and spatial and episodic memory[J]. Neuron, 2002, 35(4): 625–641.

2. Strange B A, Fletcher P C, Henson R N A, et al. Segregating the functions of human hippocampus[J]. Proceedings of the National Academy of Sciences, 1999, 96(7): 4034–4039.

3. Ho N F, Iglesias J E, Sum M Y, et al. Progression from selective to general involvement of hippocampal subfields in schizophrenia[J]. Molecular Psychiatry, 2017, 22(1): 142–152.

4. Hatanpaa K J, Raisanen J M, Herndon E, et al. Hippocampal sclerosis in dementia, epilepsy, and ischemic injury: differential vulnerability of hippocampal subfields[J]. Journal of Neuropathology & Experimental Neurology, 2014, 73(2): 136–142.

5. Bouwman M M A, Frigerio I, Lin C-P, et al. Hippocampal subfields: volume, neuropathological vulnerability and cognitive decline in Alzheimer’s and Parkinson’s disease[J]. Alzheimer’s Research & Therapy, 2025, 17(1): 121.

6. La Joie R, Perrotin A, de La Sayette V, et al. Hippocampal subfield volumetry in mild cognitive impairment, Alzheimer’s disease and semantic dementia[J]. NeuroImage: Clinical, 2013, 3: 155– 162.

7. Mueller S G, Stables L, Du A T, et al. Measurement of hippocampal subfields and age-related changes with high resolution MRI at 4T[J]. Neurobiology of Aging, 2007, 28(5): 719–726.

8. Huang Y, Coupland N J, Lebel R M, et al. Structural changes in hippocampal subfields in major depressive disorder: a high-field magnetic resonance imaging study[J]. Biological Psychiatry, 2013, 74(1): 62–68.

9. Wang Z, Neylan T C, Mueller S G, et al. Magnetic resonance imaging of hippocampal subfields in posttraumatic stress disorder[J]. Archives of General Psychiatry, 2010, 67(3): 296–303.

10. Giuliano A, Donatelli G, Cosottini M, et al. Hippocampal subfields at ultra high field MRI: an overview of segmentation and measurement methods[J]. Hippocampus, 2017, 27(5): 481–494.

11. Dumoulin S O, Fracasso A, Van Der Zwaag W, et al. Ultra-high field MRI: advancing systems neuroscience towards mesoscopic human brain function[J]. NeuroImage, 2018, 168: 345–357.

12. Moser E, Laistler E, Schmitt F, et al. Ultra-high field NMR and MRI—the role of magnet technology to increase sensitivity and specificity[J]. Frontiers in Physics, 2017, 5: 33.

13. Alexander D C, Dyrby T B, Nilsson M, et al. Imaging brain microstructure with diffusion MRI: practicality and applications[J]. NMR in Biomedicine, 2019, 32(4): e3841.

14. Novikov D S, Fieremans E, Jespersen S N, et al. Quantifying brain microstructure with diffusion MRI: theory and parameter estimation[J]. NMR in Biomedicine, 2019, 32(4): e3998.

15. Jeurissen B, Descoteaux M, Mori S, et al. Diffusion MRI fiber tractography of the brain[J]. NMR in Biomedicine, 2019, 32(4): e3785.

16. Mukherjee P, Berman J I, Chung S W, et al. Diffusion tensor MR imaging and fiber tractography: theoretic underpinnings[J]. American Journal of Neuroradiology, 2008, 29(4): 632–641.

17. Roebroeck A, Miller K L, Aggarwal M. Ex vivo diffusion MRI of the human brain: technical challenges and recent advances[J]. NMR in Biomedicine, 2019, 32(4): e3941.

18. Modo M, Sparling K, Novotny J, et al. Mapping mesoscale connectivity within the human hippocampus[J]. NeuroImage, 2023, 282: 120406.

19. Shih N-C, Kurniawan N D, Cabeen R P, et al. Microstructural mapping of dentate gyrus pathology in Alzheimer’s disease: a 16.4 Tesla MRI study[J]. NeuroImage: Clinical, 2023, 37: 103318.

20. Zhao Z, Zhang L, Luo W, et al. Layer-specific microstructural patterns of anterior hippocampus in Alzheimer’s disease with ex vivo diffusion MRI at 14.1 T[J]. Human Brain Mapping, 2022, 44(2): 458–471.

21. Oishi K, Mori S, Troncoso J C, et al. Mapping tracts in the human subthalamic area by 11.7T ex vivo diffusion tensor imaging[J]. Brain Structure and Function, 2020, 225(4): 1293–1312.

22. Ly M, Foley L, Manivannan A, et al. Mesoscale diffusion magnetic resonance imaging of the ex vivo human hippocampus[J]. Human Brain Mapping, 2020, 41(15): 4200–4218.

23. Ke J, Foley L, Hitchens K, et al. Ex vivo mesoscopic diffusion MRI correlates with seizure frequency in patients with uncontrolled mesial temporal lobe epilepsy[J]. Human Brain Mapping, 2020, 41.

24. Fritz F J, Sengupta S, Harms R L, et al. Ultra-high resolution and multi-shell diffusion MRI of intact ex vivo human brains using KT-DSTEAM at 9.4T[J]. NeuroImage, 2019, 202: 116087.

25. Beaujoin J, Palomero-Gallagher N, Boumezbeur F, et al. Post-mortem inference of the human hippocampal connectivity and microstructure using ultra-high field diffusion MRI at 11.7 T[J]. Brain Structure and Function, 2018, 223(5): 2157–2179.

26. Aggarwal M, Nauen D W, Troncoso J C, et al. Probing region-specific microstructure of human cortical areas using high angular and spatial resolution diffusion MRI[J]. NeuroImage, 2015, 105: 198–207.

27. Calabrese E, Hickey P, Hulette C, et al. Postmortem diffusion MRI of the human brainstem and thalamus for deep brain stimulator electrode localization[J]. Human Brain Mapping, 2015, 36(8): 3167–3178.

28. Aggarwal M, Zhang J, Pletnikova O, et al. Feasibility of creating a high-resolution 3D diffusion tensor imaging based atlas of the human brainstem: a case study at 11.7 T[J]. NeuroImage, 2013, 74: 117–127.

29. Seehaus A K, Roebroeck A, Chiry O, et al. Histological validation of DW-MRI tractography in human postmortem tissue[J]. Cerebral Cortex, 2013, 23(2): 442–450.

30. Kleinnijenhuis M, Zerbi V, Küsters B, et al. Layer-specific diffusion weighted imaging in human primary visual cortex in vitro[J]. Cortex, 2013, 49(9): 2569–2582.

31. Huang H, Xue R, Zhang J, et al. Anatomical characterization of human fetal brain development with diffusion tensor magnetic resonance imaging[J]. Journal of Neuroscience, 2009, 29(13): 4263–4273.

32. Shepherd T M, Özarslan E, Yachnis A T, et al. Diffusion tensor microscopy indicates the cytoarchitectural basis for diffusion anisotropy in the human hippocampus[J]. American Journal of Neuroradiology, 2007, 28(5): 958–964.

33. Coras R, Milesi G, Zucca I, et al. 7T MRI features in control human hippocampus and hippocampal sclerosis: an ex vivo study with histologic correlations[J]. Epilepsia, 2014, 55(12): 2003–2016.

34. Elston G N. Cortex, cognition and the cell: new insights into the pyramidal neuron and prefrontal function[J]. Cerebral Cortex, 2003, 13(11): 1124–1138.

35. Spruston N. Pyramidal neurons: dendritic structure and synaptic integration[J]. Nature Reviews Neuroscience, 2008, 9(3): 206–221.

36. Naidich T P, Nimchinsky E A, Pasik P. CHAPTER 10 – Cerebral cortex[M]. In: Naidich T P, Castillo M, Cha S, et al., eds. Imaging of the Brain. Philadelphia: W.B. Saunders, 2013: 154–173.

37. Amaral D G, Scharfman H E, Lavenex P. The dentate gyrus: fundamental neuroanatomical organization (dentate gyrus for dummies)[J]. Progress in Brain Research, 2007, 163: 3–22.

38. Tournier J-D, Smith R, Raffelt D, et al. MRtrix3: a fast, flexible and open software framework for medical image processing and visualisation[J]. NeuroImage, 2019, 202: 116137.

39. Tournier J-D, Calamante F, Connelly A. Robust determination of the fibre orientation distribution in diffusion MRI: non-negativity constrained super-resolved spherical deconvolution[J]. NeuroImage, 2007, 35(4): 1459–1472.

40. Descoteaux M, Angelino E, Fitzgibbons S, et al. Regularized, fast, and robust analytical q-ball imaging[J]. Magnetic Resonance in Medicine, 2007, 58(3): 497–510.

41. Jones D K. Tractography gone wild: probabilistic fibre tracking using the wild bootstrap with diffusion tensor MRI[J]. IEEE Transactions on Medical Imaging, 2008, 27(9): 1268–1274.

42. Khan A R, Cornea A, Leigland L A, et al. 3D structure tensor analysis of light microscopy data for validating diffusion MRI[J]. NeuroImage, 2015, 111: 192–203.

43. Budde M D, Frank J A. Examining brain microstructure using structure tensor analysis of histological sections[J]. NeuroImage, 2012, 63(1): 1–10.

44. Förster E, Zhao S, Frotscher M. Laminating the hippocampus[J]. Nature Reviews Neuroscience, 2006, 7(4): 259–268.

45. Calamante F, Tournier J-D, Kurniawan N D, et al. Super-resolution track-density imaging studies of mouse brain: comparison to histology[J]. NeuroImage, 2012, 59(1): 286–296.

46. Henze D A, Urban N N, Barrionuevo G. The multifarious hippocampal mossy fiber pathway: a review[J]. Neuroscience, 2000, 98(3): 407–427.

47. Geula C. Abnormalities of neural circuitry in Alzheimer’s disease[J]. Neurology, 1998, 51(Suppl 1): S18–S29.

48. Shi H-J, Wang S, Wang X-P, et al. Hippocampus: molecular, cellular, and circuit features in anxiety[J]. Neuroscience Bulletin, 2023, 39(6): 1009–1026.

49. Benes F M. Evidence for altered trisynaptic circuitry in schizophrenic hippocampus[J]. Biological Psychiatry, 1999, 46(5): 589–599.

50. Yushkevich P A, Muñoz López M, Iñiguez De Onzoño Martin M M, et al. Three-dimensional mapping of neurofibrillary tangle burden in the human medial temporal lobe[J]. Brain, 2021, 144(9): 2784–2797.

51. Mancini M, Casamitjana A, Peter L, et al. A multimodal computational pipeline for 3D histology of the human brain[J]. Scientific Reports, 2020, 10(1): 13839.

52. Adler D H, Pluta J, Kadivar S, et al. Histology-derived volumetric annotation of the human hippocampal subfields in postmortem MRI[J]. NeuroImage, 2014, 84: 505–523.

53. Augustinack J C, Helmer K, Huber K E, et al. Direct visualization of the perforant pathway in the human brain with ex vivo diffusion tensor imaging[J]. Frontiers in Human Neuroscience, 2010, 4: 42.

54. Jeurissen B, Leemans A, Tournier J-D, et al. Investigating the prevalence of complex fiber configurations in white matter tissue with diffusion magnetic resonance imaging[J]. Human Brain Mapping, 2013, 34(11): 2747–2766.

55. De Calignon A, Polydoro M, Suárez-Calvet M, et al. Propagation of tau pathology in a model of early Alzheimer’s disease[J]. Neuron, 2012, 73(4): 685–697.

56. Lewis J, Dickson D W. Propagation of tau pathology: hypotheses, discoveries, and yet unresolved questions from experimental and human brain studies[J]. Acta Neuropathologica, 2016, 131(1): 27–48.

57. Witter M P. The perforant path: projections from the entorhinal cortex to the dentate gyrus[M]// Scharfman H E, ed. Progress in Brain Research. Amsterdam: Elsevier, 2007, 163: 43–61.

58. Schilling K G, Grussu F, Ianus A, et al. Considerations and recommendations from the ismrm diffusion study group for preclinical diffusion mri: part 2—ex vivo imaging: added value and acquisition[J]. Magnetic Resonance in Medicine, 2025, 93(6): 2535–2560.

59. Zeineh M M, Palomero-Gallagher N, Axer M, et al. Direct visualization and mapping of the spatial course of fiber tracts at microscopic resolution in the human hippocampus[J]. Cerebral Cortex, 2017, 27(3): 1779–1794.

60. Oberstrass A, DeKraker J, Palomero-Gallagher N, et al. Analyzing regional organization of the human hippocampus in 3D-PLI using contrastive learning and geometric unfolding[Z/OL]. arXiv, 2024, arXiv:2402.17744.

61. Morawski M, Kirilina E, Scherf N, et al. Developing 3D microscopy with CLARITY on human brain tissue: Towards a tool for informing and validating MRI-based histology[J]. NeuroImage, 2018, 182: 417–428.

62. Xu F, Shen Y, Ding L, et al. High-throughput mapping of a whole rhesus monkey brain at micrometer resolution[J]. Nature Biotechnology, 2021, 39(12): 1521–1528.

